# An Accessible Continuous-Culture Turbidostat for Pooled Analysis of Complex Libraries

**DOI:** 10.1101/450536

**Authors:** Anna M. McGeachy, Zuriah A. Meacham, Nicholas T. Ingolia

## Abstract

We present an accessible, robust continuous-culture turbidostat system that enables biologists to generate and phenotypically analyze highly complex libraries in yeast and bacteria. Our system has many applications in genomics and systems biology; here, we demonstrate three of these uses. We first measure how the growth rate of budding yeast responds to limiting nitrogen at steady state and in a dynamically varying environment. We also demonstrate the direct selection of a diverse, genome-scale protein fusion library in liquid culture. Finally, we perform a comprehensive mutational analysis of the essential gene *RPL28* in budding yeast, mapping sequence constraints on its wild-type function and delineating the binding site of the drug cycloheximide through resistance mutations. Our system can be constructed and operated with no specialized skills or equipment and applied to study genome-wide mutant pools and diverse libraries of sequence variants under well-defined growth conditions.

## INTRODUCTION

Liquid cultures of bacteria and yeast are ubiquitous in biology research. These cultures provide the setting for diverse experiments studying microbes themselves as well as using them as a host for the expression of heterologous genes and proteins. Well-mixed liquid cultures provide a homogeneous environment for a large cell population, enabling consistent measurements of cellular physiology (1, 2). These cultures also provide a setting for experimental adaptation and evolution that can be compared easily with theoretical predictions (3, 4). Large populations grown in liquid culture also support the propagation of diverse libraries of genes or gene variants, allowing genome-wide genetic analysis and comprehensive, “deep” mutational scanning (5-7). All of these applications stand to benefit from controllable, continuous culture for steady-state growth and dynamical environmental modulations (8).

Continuous culture systems can be used to eliminate the irreproducible environmental and physiological changes that arise during batch cultures, but they require some feedback to maintain culture conditions. Chemostats establish a steady-state culture where cell growth is balanced against a fixed rate of media influx and the culture itself plays a role in the feedback loop (1, 2). Cell growth is restricted by the availability of some limiting nutrient, and cell density stabilizes at the point where consumption of this limiting nutrient matches its influx. This chemostatic steady state can occur only when cells persist in a partially starved state, and so it is best suited for certain metabolic studies.

Turbidostats maintain a steady-state culture by directly monitoring cell density and diluting the culture with fresh media to compensate for cell growth (9). Steady-state growth at a constant cell density seems conceptually simpler than chemostat growth, but it depends on active, external feedback by a controller that monitors the culture and dynamically adds media. Simple access to microcontrollers along with fabrication by 3D printing now enable individual laboratories to construct turbidostats at low cost (10-13). These systems are nonetheless complex to build and to operate, limiting their widespread adoption. Furthermore, culture sizes in these turbidostats limit the diversity of pooled libraries that they can maintain.

We present here an accessible and robust bioreactor design that maintains a defined culture condition for several days using non-invasive turbidity measurements. Our system will enable many researchers to grow large, high diversity libraries in constant conditions and aid the wider adoption of continuous culture growth. We present a simple circuit that measures cell density in ambient conditions using scattered light (nephelometry). These turbidity measurements allow us to maintain a culture at steady state while determining its growth rate. By algorithmically adjusting media composition, we are able to map the response of the budding yeast *Saccharomyces cerevisiae* to nitrogen limitation and drive the culture through reproducible cycles of feast and famine. Our bioreactor can support populations of over 10^9^ cells, and we show how this enables the construction and phenotypic analysis of complex transformant pools directly in liquid culture. Using complex selective regimes enabled by the turbidostat, we perform deep mutational scanning on the gene *RPL28*, which encodes an essential ribosomal protein (14) that defines, in part, the binding site for the drug cycloheximide (15). Our results profile comprehensively the effects of amino acid substitutions on normal Rpl28 function. They also reveal alleles conferring resistance to the ribosomal poison cycloheximide, which map the known binding site of this drug on the ribosome. The results we present here exemplify the diverse experiments that are enabled by this approachable and robust culture system. In order to support the broad adoption of this approach in systems and synthetic biology, we provide design files and detailed instructions that allow other laboratories to easily build and use this system (see https://github.com/ingolia-lab/turbidostat).

## RESULTS

### Cell density measurement by scattered light

Turbidity serves as an excellent proxy for cell density in liquid culture. While culture density is routinely measured by light absorbance, many challenges arise in measuring absorbance in bioreactors. We found that scattered light provided a robust measure of turbidity across a wide dynamic range. Our turbidity detector features a near-infrared LED illuminating the culture and a photodetector placed at a 90° angle, with the geometry maintained by a custom plastic housing (Fig. 1a). By modulating the illumination at 10 kHz, we extract the faint scattered light signal out of the ambient background using synchronous detection combined with electronic filters tuned to this frequency (Supplementary Fig. 1) (16). Light modulation and detection are accomplished by an inexpensive microcontroller that can also control the addition of media to the culture.

Our nephelometer can measure the range of cell densities commonly used in microbial culture. Scattered light is clearly detectable above background in cultures of the budding yeast *S. cerevisiae* (Fig. 1b and Supplementary Fig. 2a and 2b) and the bacterium *E. coli* (Fig. 1c and Supplementary Fig. 2c and 2d) at OD600 as low as 0.05, and the intensity scales linearly with cell density up to an OD600 of 1.0. Signal intensity continues to increase beyond this point, and so the system is suitable for continuous culture of higher cell densities as well.

### Feedback control for continuous culture

We designed a continuous culture bioreactor using readily available components that is suitable for autoclave sterilization after assembly. The growth chamber, a standard 250 ml or 500 ml glass bottle, is maintained at positive pressure by the continuous flow of sterile, hydrated air (or other gas) (Fig. 1d and 1e). Media is added into the growth chamber by a peristaltic pump, and excess culture volume is displaced out of the waste line by positive pressure in the growth chamber. Our design supports two pumps for two different media sources, driven by the microcontroller that monitors cell density, and it could easily accommodate additional pumps or sensors. The fluidics uses common laboratory glassware connected by silicone tubing and assembled with luer fitting, and requires no custom components (Fig. 1d).

We established a steady-state *E. coli* culture by linking the addition of fresh media to cell density measurements, implementing a turbidostat. Stationary-phase *E. coli* entered exponential growth after ∼10 hours and cell density increased exponentially until cells reached our target turbidity (Fig. 1f). From this point, the influx of fresh media maintained a constant turbidity, and the rate of media addition stabilized at a dilution rate of 0.39 hr^-1^ (Fig. 1f); as dilution matches growth when cell density is constant, we inferred a doubling time of 1.8 hours, consistent with our expectation for *E. coli* grown at ∼20 °C.

**Figure 1.**
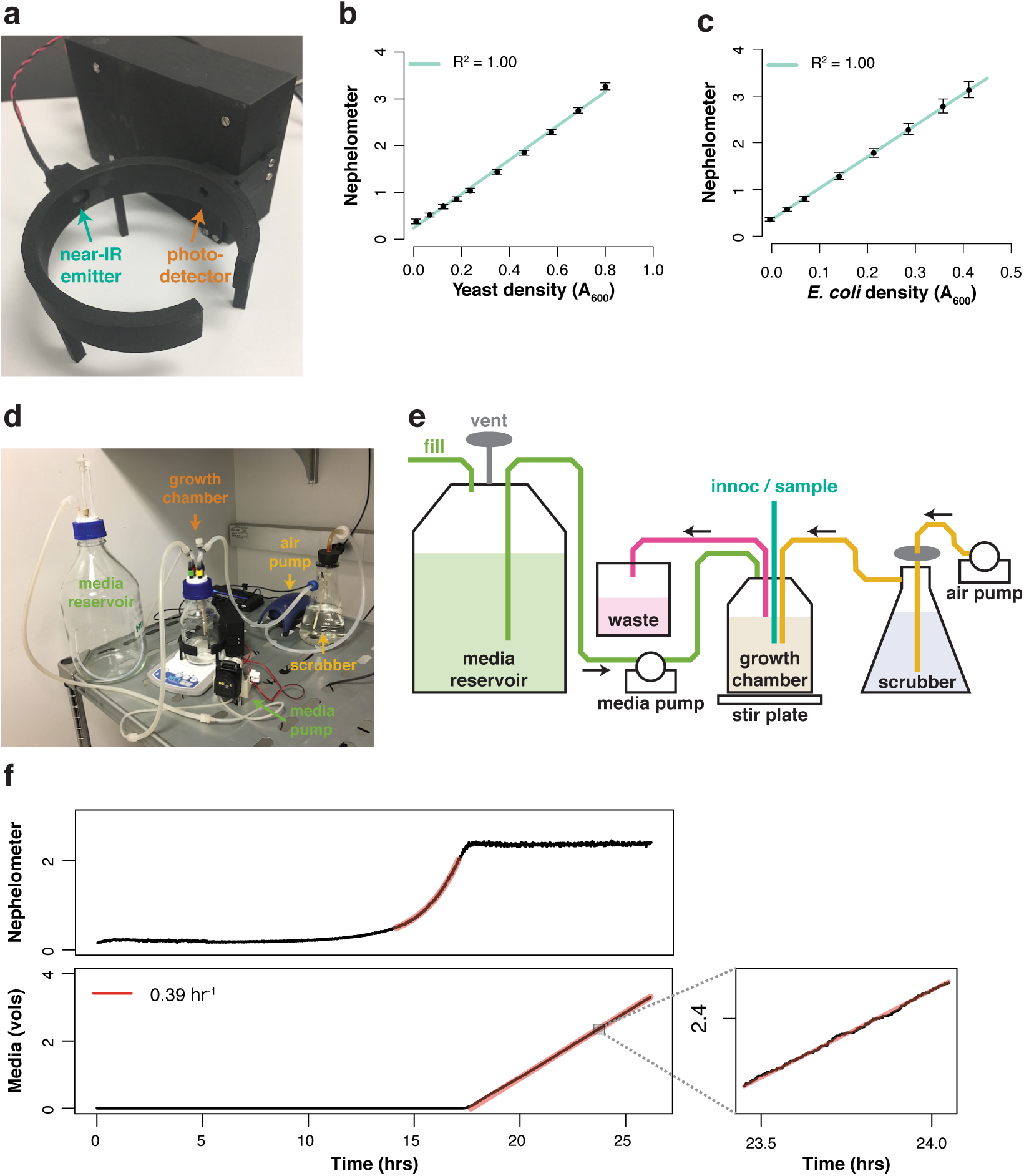
Turbidostatic continuous culture of bacteria and yeast. (**a**) Turbidity detector for cultures in a 250 ml growth chamber. (**b, c**) Scattered light turbidity (nephelometry) of yeast and bacterial provide precise, linear measurements of cell density. (**d, e**) Continuous culture growth system. A photograph of the assembled system is shown in (d), and a schematic of the fluidics is shown in (e). (**f**) Continuous culture of bacteria maintained in a turbidostat. The rate of media addition and culture dilution needed to maintain constant cell density is uniform and reflects the growth rate.

### Cellular physiology in dynamic environments

Our system can control two different media sources simultaneously, allowing us to produce complex and dynamic environments. We used this feature to maintain budding yeast cultures at progressively lower concentrations of ammonium and measure how nitrogen availability affected growth rate. Our turbidostat algorithm estimates the current media composition of the growth chamber and, when media addition is required, chooses between minimal media with and without a nitrogen source in order to quickly reach and then maintain a target nitrogen concentration. Yeast minimal media typically contains 76 mM NH_4_^+^ (17), and in preliminary experiments we determined that the growth rate of prototrophic S288c haploids remained unchanged at NH_4_^+^ levels as low as 2.0 mM (Supplementary Fig. 4). We then followed the growth of yeast grown with NH_4_^+^ levels declining sequentially from 4.0 mM until growth ceased (Fig. 2a). Growth rate appeared to stabilize ∼3 hours after a stepwise decrease in ammonium concentration (Fig. 2b), and so we conservatively allowed the culture to equilibrate at each concentration for 6 hours prior to calculating an average growth rate over the final 2 hours.

Our data revealed a modest growth decrease below 2.0 mM NH_4_^+^, prior to entering a tightly nitrogen-limited regime around 500 μM NH_4_^+^ (Fig. 2c). The 2.0 mM threshold for reduced growth matched the estimated KM of the low-affinity, high-capacity Mep3p ammonium transporter (18). We fit the linear relationship between growth rate and ammonium content in the nitrogen-limited regime and extrapolated that growth ceases at a nitrogen concentration of ∼120 μM (Fig. 2c). This value agreed with the concentration of fixed nitrogen we expected in our culture, grown at 2.5e6 cells / ml, showing that growth stopped when incoming free nitrogen dropped below the amount lost in outgoing yeast (19). We then fixed the media composition at 400 μM NH_4_^+^ and varied cell density, again finding a linear decrease in growth rate that extrapolated to zero at 7.7e6 cells / ml, where the concentration of fixed nitrogen in the culture matches the concentration of nitrogen in the incoming media. Growth under these low-nitrogen conditions is similar to chemostat culture, though we are able to vary cell density and media composition as two independent variables, whereas only the dilution rate can be controlled in a chemostat. Turbidostat culture allows us to establish cultures at a steady state determined more by experimental design and affected less by dynamic responses of the culture.

Our turbidostat also allows us to vary the composition of our medium during growth and create a changing culture environment. We varied the nitrogen composition of our medium between 300 μM ammonium, which severely restricts growth, and 2.0 mM ammonium, which does not appear to be limiting (Fig. 2f). Growth rates varied roughly 3-fold across the 16-hour cycle, and depended strongly on the history of the culture (Fig. 2g). Cells quickly stopped dividing when ammonium levels dropped below ∼600 μM, but they were slower to resume growth as nitrogen availability increased. In fact, growth rate continued to rise for ∼4 hours after the nitrogen concentration peaked.

**Figure 2.**
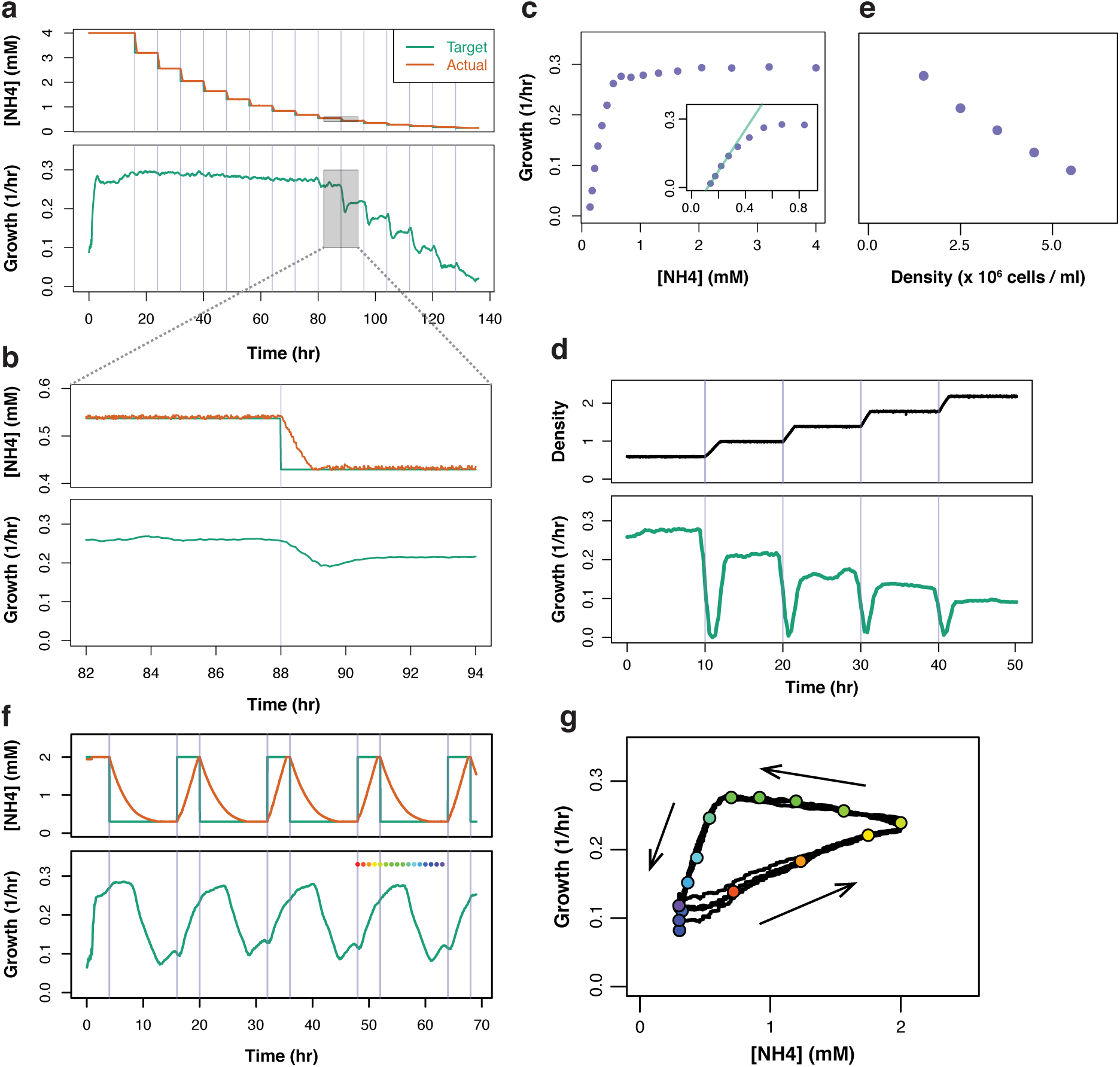
Measuring the response of yeast culture to varying nitrogen levels. (**a**) Turbidostat culture maintained at progressively lower [NH4+] levels. (**b**) Growth drops transiently after nitrogen reduction before equilibrating to a new, slower rate. (**c**) Changes in steady-state growth rate as a function of [NH4+] levels in the medium. Inset shows a magnification of the response at low [NH4+] levels, along with a linear fit in the strongly nitrogen limited regime. (**d**) Turbidostat culture in limiting nitrogen (400 μM NH_4_^+^) maintained at progressively higher cell density. (**e**) Changes in the rate of steady-state growth in 400 μM NH_4_^+^ at varying cell density. (**f**) Periodic changes in media composition drive cyclic changes in cell growth rate. Rainbow dots indicate 1-hour timepoints across one cycle of levels. (**g**) Phase plot of yeast culture growth responding to periodic changes in nitrogen availability. Rainbow dots correspond to timepoints in (f).

### Pooled selective growth of diverse libraries

Our growth chamber supports populations of billions of cells, enabling the pooled growth and phenotypic analysis of complex libraries. These complex libraries can support a wide range of experiments: large pools of mutants for forward genetic analysis, comprehensive variant libraries for deep mutational scanning, or proteome-wide protein fusion libraries for functional experiments such as yeast two-hybrid analysis.

We first tested our ability to select an expression library of in-frame protein fusions. Expression libraries constructed directly from genomic DNA (gDNA) or from whole-transcriptome complementary DNA (cDNA) suffer from a large fraction of non-coding and out-of-frame fragments. It is possible to select for in-frame fusion proteins by expressing the selectable marker gene as a translational fusion downstream of the captured gDNA or cDNA fragment (20). In this configuration, expression of the selectable marker depends on the correct translation of the upstream fusion protein. This approach has not been widely adopted, however, perhaps because translational fusion can disrupt either the selectable marker or the library fragment itself. We noted that multi-cistronic expression of two proteins from a single reading frame can now be achieved through the use of viral “self-cleaving” 2A peptides (21). These short peptides autonomously skip one peptide bond during translation by eukaryotic ribosomes, producing two separate proteins (22).

We took advantage of 2A peptides to design a vector for recombinational cloning and selection of an inframe expression library in yeast. Recombinational cloning allows us to capture a diverse array of gDNA or cDNA fragments into our vector by adding short, 40 base pair homology arms on each end of the fragment and co-transforming them along with the vector into budding yeast, where homologous recombination will efficiently assemble a circular plasmid (23). The vector contains an auxotrophic yeast marker with a 2A peptide, but it is not expressed unless a captured fragment reconstitutes an intact reading frame upstream of the marker (Fig. 3a).

In order to produce a diverse pool of input DNA fragments with minimal bias, we adopted the approaches used to construct high-throughput sequencing libraries, which have been optimized with the goal of complete and uniform coverage of input DNA (24). These sequencing libraries also feature known adapters at each end of the fragment. We incorporated a part of those adapters into our vector, allowing us to amplify templates for recombinational cloning directly from a sequencing library.

We generated a diverse expression library from yeast genomic DNA and analyzed its composition by deep sequencing. Our strategy incorporates a portion of sequencing adapters directly into the vector, and so we were able to easily amplify a high-throughput sequencing library from these plasmids. Prior to selection, most DNA fragments that we sequenced contained in-frame stop codons and we saw no preference for any particular codon phase in fragment length (Fig. 3b). After selection, however, over half of the DNA fragments we sequenced are expected to produce an intact fusion protein — they lack stop codons and establish the correct reading frame across the insertion point (Fig. 3b). Indeed, these fragments were strongly enriched during selection, at the expense of those that disrupted the reading frame (Fig. 3c).

Selection for frame-preserving insertions captured a diverse pool of yeast protein fragments. We saw enrichment only for insertions that placed yeast coding sequences (CDSes) in the correct reading frame relative to our expression vector (Fig. 3d). Fragments in other reading frames, as well as those that fell outside a CDS, were depleted during selection. Our library contained multiple fragments from most yeast genes, and gene length was a major determinant of the number of fragments that we saw (*r* = 0.64) (Fig. 3e). Individual fragments were typically large enough to contain a well-folded structural motif (Fig. 3f), and the library size could presumably be controlled by changing the size distribution of the input DNA. Based on these results, we conclude that turbidostat selection is a valuable approach for generating diverse expression libraries.

**Figure 3.**
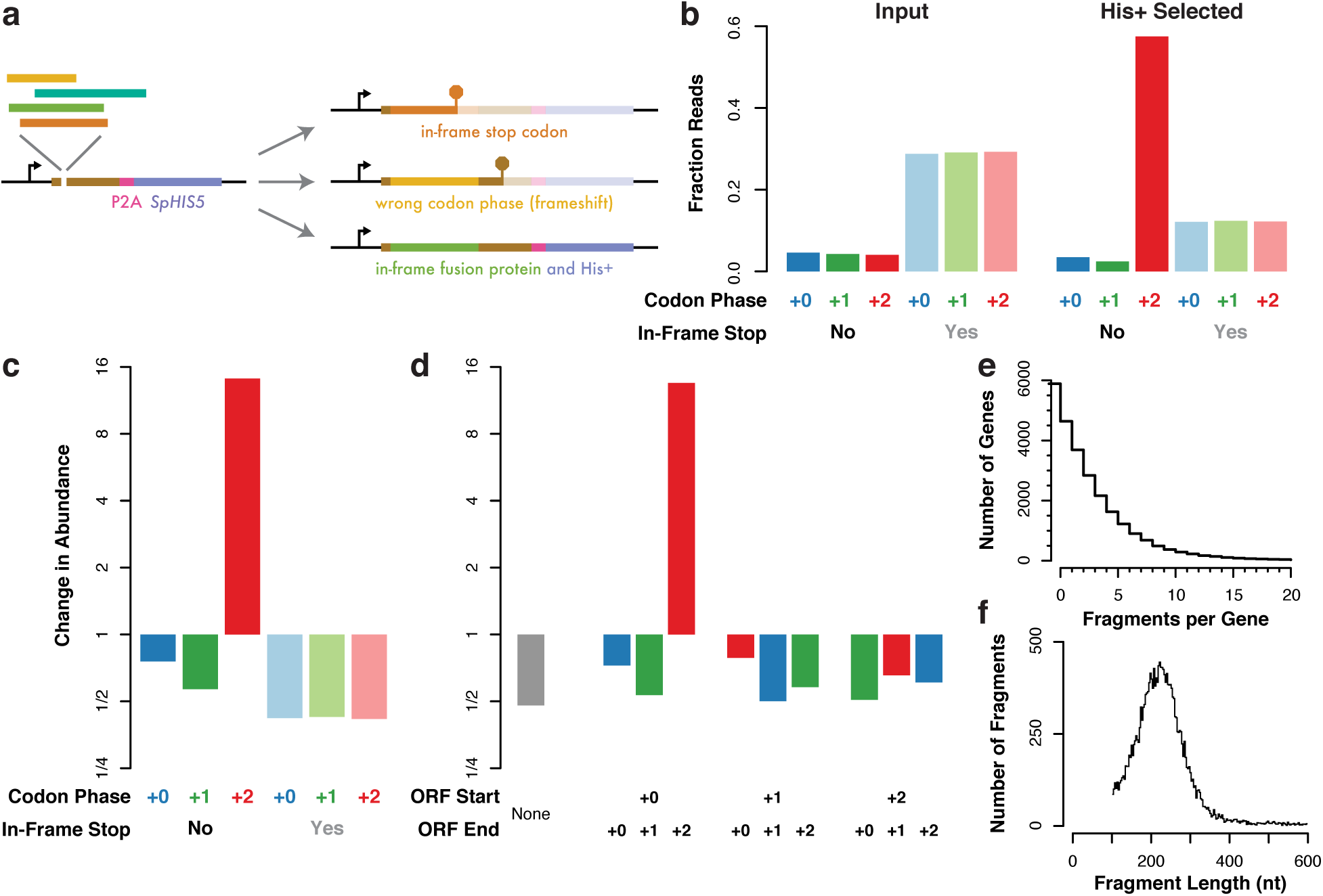
Constructing a proteome-wide in-frame expression library. **(a)** Expression of the downstream selectable marker depends on in-frame translation of an upstream library fragment. **(b)** Reading frame distribution across a library of yeast genomic DNA fragments. Prior to selection, fragments typically contain stop codons and show no codon phase bias. After selection, most fragments lack stop codons and produce a +2 reading frame across the insertion site, which is compatible with inframe translation of the downstream marker. **(c)** Specific enrichment of frame-preserving fragments out of a yeast genomic DNA library after selection for the downstream marker. **(d)** Enrichment of genomic DNA fragments derived from annotated protein-coding genes and translated in the correct reading frame. Codon phases are colored as in (b) and (c). **(e)** Cumulative distribution of the number of in-frame fragments per yeast gene. **(f)** Length distribution of fragments.

### Phenotypic characterization of diverse libraries in continuous culture

Having established that we could select diverse libraries directly in continuous culture, we next wanted to characterize the growth phenotypes of these transformed populations. We set out to characterize variants of *RPL28*, an essential, single-copy gene encoding a ribosomal protein, the yeast uL15 orthologue (14). The translation inhibitor cycloheximide binds the yeast ribosome in a pocket defined by Rpl28p along with Rpl44p and the 28S ribosomal RNA (15), and a glutamine to glutamate mutation at position 38 of *RPL28* confers cycloheximide resistance (25). We wished to scan the phenotypes of *RPL28* variants to determine which alleles were viable, and to identify the full range of mutations conferring cycloheximide resistance. As *RPL28* is essential, we needed to provide wild-type *RPL28* function and then remove the wild-type gene to expose the phenotype of our mutants. We thus complemented a deletion of genomic *RPL28* with an episomal plasmid containing the counter-selectable yeast marker *URA3*. This strain can be used to perform a two-step plasmid shuffle, in which a second episomal plasmid is introduced, and cells carrying the *URA3*-marked wild-type plasmid are eliminated by treatment with the toxic uracil analogue 5-FOA (Fig. 4a) (26).

We implemented this entire genetic scheme on a library of ∼270,000 distinctly barcoded plasmids in continuous turbidostat culture.We transformed this library into yeast and inoculated the transformation directly into our turbidostat in order to select for His+ transformants. This transformation yielded ∼7.5 million transformants (>25x coverage of the library) from ∼800 million total yeast using 24 μg of library DNA. Substantial growth began ∼12 hours after inoculation (Fig. 4b), and after 24 hours we found that ∼15% of cells lacked the wild-type plasmid and ∼31% of cells lacked a variant plasmid, though we isolated no clones lacking both plasmids (Fig. 4c). We attribute the high fraction of cells with no variant library plasmid (Ura+ His-) to the perdurance of the His3 protein product in cells that lose the plasmid spontaneously, combined with incomplete dilution of untransformed cells out of the culture. In contrast, the simultaneous loss of both *RPL28* plasmids stops growth almost immediately, and so we recover no Ura‐ His‐ cells.

We then added 5-FOA to our media in order to eliminate cells with wild-type *RPL28* and expose the phenotypes of our variant library. Addition of 5-FOA depressed growth for ∼8 hours before the culture recovered and grew robustly (Fig. 4b). All clones isolated after ∼22 hours of 5-FOA counter-selection lacked the *URA3*-marked wild-type plasmid (Fig. 4c). After eliminating wild-type *RPL28*, we switched our turbidostat to media with 200 nM cycloheximide and no 5-FOA. Cycloheximide quickly and dramatically suppressed growth, reaching a plateau below half of the initial growth rate before recovering slightly (Fig. 4b).

We assessed the growth phenotype of individual *RPL28* variants across this selection scheme by tracking the abundances of the associated barcodes. We amplified these barcodes from yeast sampled at six different timepoints (Fig. 4b), as well as the input DNA library used to transform yeast, and quantified them by deep sequencing. Barcode abundance in transformed yeast correlated very well at different timepoints, and also correlated with the composition of the input DNA library (Fig. 4d and 4e). These abundances diverged substantially after selection was applied, however, reflecting the different growth phenotypes of the linked *RPL28* variants (Fig. 4f). We clustered samples by barcode abundance and found three distinct groups with clear phenotypic meanings. One group reflected the initial plasmid library, another comprised the protein variants that support growth in the absence of wild-type *RPL28*, and the last showed dramatic enrichment of alleles conferring cycloheximide resistance.

**Figure 4.**
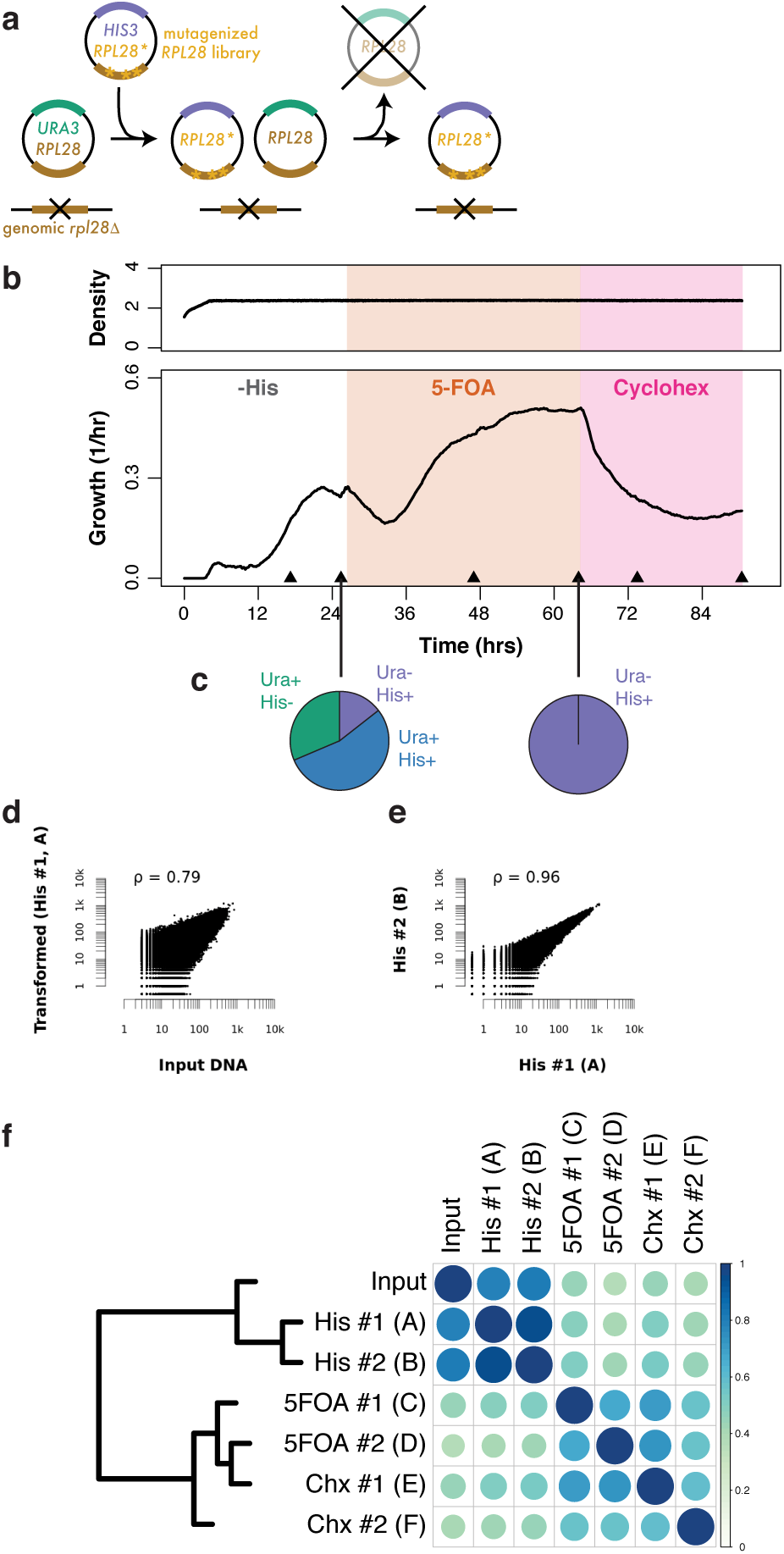
Growth of an *RPL28* variant library in continuous culture. (**a**) Plasmid exchange strategy to introduce *RPL28* variants and then eliminate wild-type *RPL28*. (**b**) Turbidostat culture of yeast during transformation of a barcoded variant library, counter-selection against the wild-type *RPL28* plasmid, and cycloheximide treatment of yeast subsisting on *RPL28* variants. (**c**) Frequency of yeast containing wild-type (Ura+) and/or variant (His+) *RPL28* plasmids during selection. (**d**) Good correlation between the abundance of barcodes in the input plasmid library and the transformed yeast population assessed by deep sequencing. (**e**) Excellent correlation between the abundance of barcodes in the transformed yeast population sampled at two different timepoints. (**f**) Correlations between barcode abundances in each sample. Barcode abundances change dramatically after counter-selection against wild-type *RPL28* and change further during growth in 200 nM cycloheximide.

### Fitness effects of *RPL28* mutations

In order to interpret our phenotypic data, we determined the *RPL28* sequences associated with the barcodes in our variant library. We generated a high-throughput sequencing library with barcodes on one end and random breakpoints in the *RPL28* gene on the other end. Deep sequencing of this library provided a barcode sequence linked to a region of *RPL28* sequence in a mated sequence pair (27). As the mutagenized region of *RPL28* was ∼700 base pairs long, many distinct fragments were needed to achieve full sequence coverage, and we only captured variant information for a fraction of our barcodes. We focused on these barcodes where we knew the complete *RPL28* sequence when analyzing our fitness data.

Growth after 5-FOA selection reflected the fitness of individual *RPL28* alleles. Individual barcodes showed a bimodal fitness distribution, with one class of barcodes dropping ∼30-fold in abundance relative to the other (Fig. 5a). Overall, barcodes associated with wild-type *RPL28* alleles fell into the high-fitness group while those associated with presumptively non-functional alleles containing nonsense or splice acceptor site mutations had low fitness. We then found the fitness of 665 single-nucleotide variants by collecting all distinct barcodes linked to an *RPL28* allele bearing only that mutation. Synonymous and intronic mutations, which are expected to be phenotypically silent, indeed displayed uniformly high fitness (Fig. 5b). In contrast, mutations that disrupt the *RPL28* reading frame showed consistently low fitness. Based on these easily understood mutations, we defined thresholds that divided our fitness landscape into three broad classes, “healthy”, “sick”, and “dead” (Fig. 5b). Individual mis-sense mutations spanned the full range of fitnesses from silent to lethal.

Looking at these mis-sense mutations, we found that the fitness of amino acid substitutions tracked the severity of the change. We grouped mis-sense variants by amino acid substitution and found that sick and lethal alleles were substantially enriched for disfavored changes with low scores in the PAM30 substitution matrix (Fig. 5c) (28). We then took advantage of the deep evolutionary conservation of *RPL28* and tested substitutions at each amino acid position against the position-specific scoring matrix (PSSM) built across a wide range of eukaryotic orthologues (CHL00112) (29). More deleterious substitutions were more likely to occur at positions where the budding yeast wild-type residue scored highly in the PSSM and the mutant residue scored poorly (Fig. 5d). Together, these effects produced a strong correlation (ρ = 0.44) between the fitness effect of a mutation and the wild-type to mutant PSSM score difference — a stronger effect than the correlation with the generic PAM30 score (ρ = 0.31).

**Figure 5.**
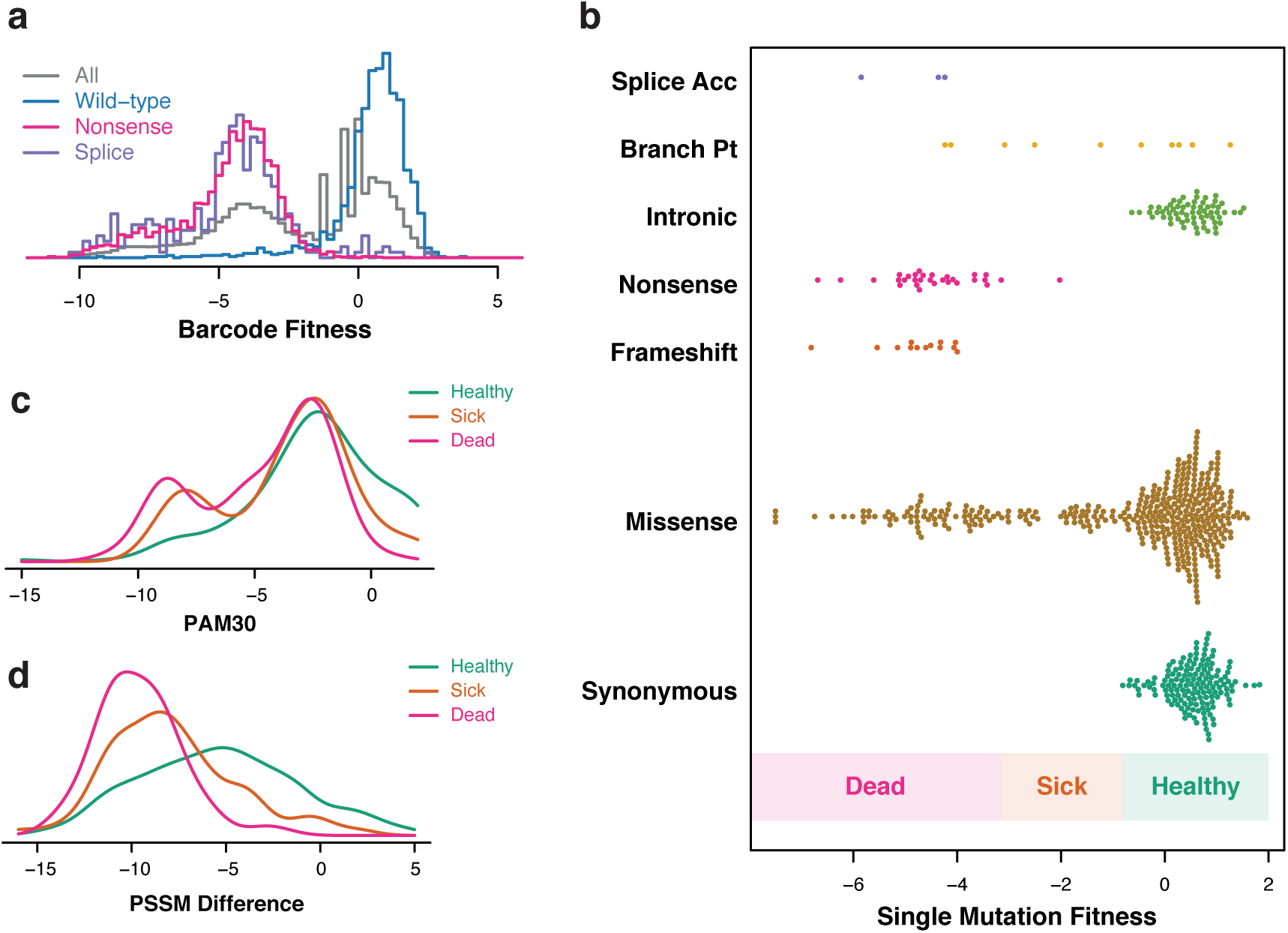
Phenotypic characterization of *RPL28* variants. (**a**) Histogram of fitness measurements for individual barcodes associated with wild-type *RPL28* and with presumptive null alleles. (**b**) Fitness effects of single nucleotide mutants categorized by their effect on *RPL28*. The threshold for “healthy” fitness is the lowest measurement among synonymous mutations while the threshold for “dead” fitness is the highest measurement among presumptive null mutations. (**c**) Density plot for amino acid substitution score in the PAM30 matrix for missense mutations in the “healthy”, “sick”, and “dead” fitness class. (**d**) Density plot as in (c) for the difference in PSSM score between the wild-type and mutant residue.

### Cycloheximide resistance alleles delineate its binding site on the ribosome

We next tracked how the abundance of viable *RPL28* alleles changed during growth in cycloheximide. Sick alleles continued to show a fitness defect, whereas the overall distribution of healthy mutants was similar to that of wild-type barcodes (Fig. 6a). Notably, a small number of healthy mutant barcodes showed a strong fitness advantage in cycloheximide, whereas wild-type barcodes showed no such effect. We then examined the fitness of amino acid substitutions and found two regions where mutations enhanced fitness: Gly37/Gln38/His39 and Gly54/Lys55/Arg59 (Fig. 6b). Notably, these mutations included the Gln38Glu substitution underlying the “*cyhx-32*” cycloheximide resistance allele (25), as well as all other substitutions observed at this position (to His, Leu, Lys, Pro, and Arg). We selected a chemically distinct substitution, Gln38Pro, for targeted validation. We also tested His39Leu, the most non-conservative resistance mutation we found (PAM30 = −6, PSSM difference = −13), along with Lys55Met, the strongest allele in the second cluster of amino acids. Each of these three mutations provided substantial cycloheximide resistance relative to wild-type *RPL28*, validating the accuracy of our global analysis (Fig. 6c and Supplementary Fig. 4).

Cycloheximide does not bind directly to Rpl28 on the ribosome. Instead, it binds a pocket comprised of ribosomal RNA and the protein Rpl41, and Rpl28 residues interact with the rRNA bases that contact the drug directly (15). Our strong cycloheximide resistance alleles map precisely to the residues forming this second shell around the binding site, and weaker alleles change residues that could be involved more distantly in organizing the cycloheximide pocket (Fig. 6d). The remarkable correspondence between our fitness measurements and the physical structure emphasize the value of our continuous culture system, which enables comprehensive and high resolution phenotypic analysis.

**Figure 6.**
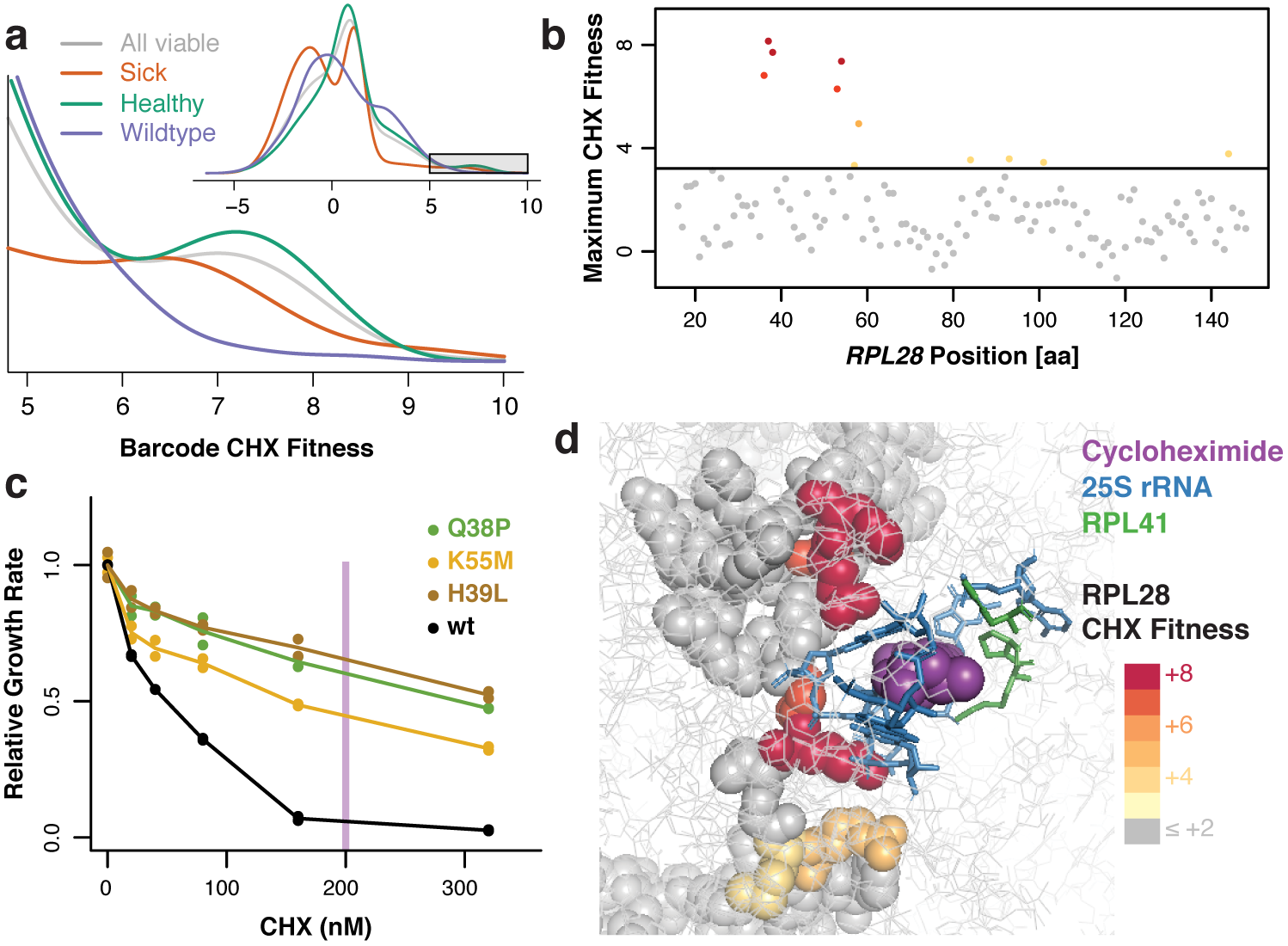
Cycloheximide resistance alleles in *RPL28*. (**a**) Density plot of fitnesses of individual barcodes. The overall distribution of viable barcodes is shown along with the distribution for sick barcodes, healthy barcodes, and barcodes linked to wild-type *RPL28*. The full density plot is shown as an inset. (**b**) Highest fitness effect seen for single residue substitutions at each position in *RPL28*. The threshold for resistance is set according to the highest fitness measured for synonymous single base changes. (**c**) Growth of individual *RPL28* variants at different cycloheximide concentrations confirms the resistance phenotype of variants identified by sequencing. (**d**) Structure of cycloheximide (purple) bound to the yeast 80S ribosome. The immediate binding pocket comprises ribosomal RNA (blue) and Rpl41 (green). Rpl28 is shown with spheres and colored according to the highest fitness observed for single residue changes as in (b).

## DISCUSSION

We have demonstrated a simple and accessible turbidostat with many uses in systems biology. Our system can measure growth and control dynamically varying culture conditions, enabling studies of cell physiology. Relative to other continuous culture systems, construction of our turbidostat is approachable for researchers with limited engineering experience and at modest cost. The fluidics in our turbidostat are composed from common, fairly inexpensive laboratory equipment. Custom parts can be produced by many vendors, and the circuits can be assembled in a straightforward manner by following clear and simple instructions. The system is controlled by open-source software written in the “Arduino” environment and data is logged in real time over an ordinary USB connection to a computer. These design decisions ensure that the turbidostat is adaptable and robust as well as being easy to build.

One distinctive feature of our system is the larger culture volume (200 ‐ 400 ml), which enables growth of large populations (several billion cells) for the analysis of diverse libraries. We were thus able to perform complex phenotypic characterization of a comprehensive library of mutants in an essential gene, mapping the general requirements for its normal function as well as the specific residues conferring drug resistance. Pooled analysis of yeast libraries has immediate applications in studying the molecular genetics of yeast itself (6). Yeast also serves as a host for other experiments, including many approaches to characterize human proteins and their interactions (5), and more generally as a platform for synthetic biology (30) and protein engineering (31). Our turbidostat thus promises advantages in a wide array of high-throughput experiments.

## METHODS

### Turbidostat construction

Design files and detailed instructions for turbidostat construction are available at https://github.com/ingolia-lab/turbidostat.

### Calibrating turbidity and cell density

#### Figure 1b and 1c.

The correlation between turbidity, optical density (A_600_), and cell density was determined by preparing a growth chamber filled with media, aerated, and stirred. Known volumes of cells from a saturated culture were added, cells were allowed to mix evenly into the growth chamber by stirring for >100 seconds, and turbidity was measured each 0.5 seconds for 100 seconds. Samples of culture were taken at certain points for absorbance measurements of cell density. Yeast experiments used a saturated culture of *S. cerevisiae* strain BY4741 (ThermoFisher) grown 36 hours at 30 °C in YEPD with vigorous shaking. Bacterial experiments used a saturated culture of *E. coli* DH5α grown 36 hours at 37 °C in LB with vigorous shaking.

### Bacterial turbidostatic growth

#### Figure 1f.

Bacterial turbidostatic growth (Fig. 1f) was established by preparing a sterilized growth chamber and sterilized media reservoir filled with LB. The growth chamber was inoculated with 1 ml of a saturated culture of *E. coli* DH5α (NEB), resulting in an A_600_ ∼ 0.3. The culture was grown with aeration and stirring for ∼26 hours, with turbidity measurements taken every second. Media was controlled in order to maintain turbidity at an A_600_ ∼ 5.0.

### Yeast turbidostatic growth in limiting nitrogen

#### Figure 2a through 2g.

Prototrophic S288c yeast were generated by transforming BY4741 yeast with pHLUM v2 (pNTI599) (32), which complements all auxotrophic markers in this strain, by standard lithium acetate transformation. An overnight culture of prototrophic S288c was grown in 10 ml of synthetic complete drop-out (SCD) ‐His, to an A_600_ ∼ 6.6, and 7.5 ml of this culture was used to inoculate a ∼225 ml culture at a density of A_600_ ∼ 0.25.

A 10x stock of minimal media without nitrogen was prepared using 10x yeast nitrogen base without amino acids and ammonium sulfate (YNB; 15.3 grams / liter, BD 233510) along with 200 g / liter dextrose. Minimal media without nitrogen was prepared by diluting this stock 10-fold in water. Minimal media with 4 mM nitrogen was prepared in the same way, with the addition of 1 ml per liter of a 2M ammonium sulfate stock (2.64 g (NH4)2SO4 per liter), noting that this provides 2 molar equivalents of nitrogen. Media was filter sterilized through a Sterivex filter (EMD Millipore SVGPL10RC) into a sterile media reservoir.

#### Figure 2a through 2c.

A sterilized 250 ml growth chamber was pre-filled with 225 ml media with 4 mM nitrogen and inoculated. This culture was grown for 16 hours at 30 °C with aeration and stirring, and media with 4 mM nitrogen was added to maintain a turbidity roughly 10% over the initial inoculation turbidity, which corresponds to an A_600_ ∼ 0.25. The target nitrogen level of the culture was then reduced to 80% of the current value every 8 hours. The choice between media with 4 mM nitrogen and no nitrogen was controlled by maintaining a current estimate of the nitrogen in the growth chamber media, updating this estimate as new media was added, and selecting the media that would bring the nitrogen levels closer to the target. Growth rates were estimated from the rate of media addition over the final 2 hours of an 8-hour increment.

#### Figure 2c and 2d.

The ratio of media addition was fixed to yield 0.4 mM nitrogen (10% media with 4 mM nitrogen and 90% media with no nitrogen). The target density was reduced to 60% of the initial target value (corresponding to an A_600_ ∼ 0.15), and then increased by 40% of the initial target value (corresponding to increments of A_600_ ∼ +0.1) every 10 hours.

#### Figure 2f and 2g.

The turbidostat was reconfigured in order to target 2 mM nitrogen for 4 hours, followed by 0.3 mM nitrogen for 12 hours.

### Constructing an expression library of fusion proteins

#### Genomic fragment library.

Yeast genomic DNA was fragmented by sonication on a Covaris S220 sonicator at 5% duty factor, 175W peak incidence power, 200 cycles per burst, and 25 seconds. DNA fragments were purified with AMpure XP beads (Beckman Coulter A63881) at a ratio of 0.8 bead volumes per 1 sample volume according to the manufacturer’s protocol. The NEBNext Ultra DNA Library Prep Kit (NEB E7370S) was used to convert 330 ng of purified DNA fragments into a library according to the manufacturer’s protocol, stopping before the “PCR Amplification” step. The pre-selection library was amplified according to the manufacturer’s “PCR Amplification” protocol, except that only 2.0 μl of adapter ligated DNA fragments were used as template, custom primer NI-NI-38 was used in place of the index primer, and 15 cycles of amplification were performed. The pool of fragments for recombinational cloning was produced in an 800 μl PCR using Q5 polymerase (NEB M0491L) according to the manufacturer’s protocol, using 2.0 μl of adapter ligated DNA fragments as template along with primers NI-AM-197 and NI-AM-199 for 21 cycles of amplification with a 65 °C annealing temperature. Amplified DNA was purified using 400 μl (0.5 volumes) of AMpure XP beads according to the manufacturer’s protocol, with two sequential elutions in 30.0 μl of 10 mM Tris, pH 8.0, yielding 5.0 μg of DNA in total.

#### Expression library vector.

The vector for recombinational cloning was produced by digesting 20 μg of pNTI347 in a 400 μl reaction using 20.0 μl of AscI (10 U / μl; NEB R0558L) according to the manufacturer’s protocol and purified using a DNA Clean & Concentrator-25 (Zymo Research D4005) according to the manufacturer’s instructions.

#### Recombinational cloning and selection.

Vector and insert DNA were co-transformed into diploid S288c yeast by standard lithium acetate transformation at a 40x scale. The transformation was inoculated into a turbidostat and grown in SCD ‐His for 42 hours. Yeast were harvested from the turbidostat by filtration, resuspended in 40 ml 1x phosphate-buffered saline (PBS), and fixed by adding 15 ml of 16% paraformaldehyde (Electron Microscopy Science 15710) and incubating 30 minutes at room temperature in the dark. Cells were washed twice in 1x PBS and stored in 1x PBS at 4 °C.

#### Recombinant plasmid recovery.

DNA was recovered from an aliquot of fixed cells by pelleting cells and resuspending in 20 μl Y-Lysis buffer (Zymo Research Y1002-1-6) with 1.0 μl Zymolyase (Zymo Research E1004) and incubating 15 minutes at 37 °C. 1.0 μl of proteinase K was added and incubated 1 hour at 60 °C. DNA was extracted by adding180 μl water, followed by 200 μl (1 volume) chloroform, and vortexing cells for 2 minutes at room temperature. The mixture was centrifuged 10 minutes at 20,000 x g at room temperature and the upper, aqueous phase was recovered. DNA was precipitated from the aqueous phase by adding 1.0 μl GlycoBlue (ThermoFisher AM9515), 15 μl of 5M NaCl and 400 μl 100% ethanol, chilling 30 minutes at −20 °C, and centrifuging 30 minutes at 20,000 x g, 4 °C. The supernatant was removed and the DNA pellet was washed in 0.5 ml ice-cold 70% ethanol and then air dried. The extracted DNA was then resuspended in 10.0 μl of 10 mM Tris, pH 8.0.

#### Sequencing library construction.

A sequencing library was amplified from extracted DNA in two sequential 50 μl PCR reactions using Q5 polymerase according to the manufacturer’s protocol. The first reaction used 2.0 μl extracted DNA as a template with primers NI-AM-226 and NI-AM-227, and 10 cycles of amplification were performed in a two-step reaction (72 °C annealing temperature) with a 90 second annealing/extension step. The second reaction used 10.0 μl unpurified first-round PCR as a template with primers NI-AM-228 and NI-NI-3, and 10 cycles of amplification were performed in a two-step reaction (72 °C annealing temperature) with a 90 second annealing/extension step. The product was purified using 35 μl (0.7 volumes) of AMpure XP beads.

#### Deep sequencing and analysis.

Libraries were sequenced on an Illumina HiSeq 2500 using paired-end sequencing with 100 bases per read. Paired-end sequencing reads were aligned to the *S. cerevisiae* genome using Bowtie2 (33) with a maximum insert size of 1200 base pairs. The reading frame content of these fragments was analyzed using SAMtools (34) and BEDtools (35) along with custom scripts available at https://github.com/ingolia-lab/turbidostat/tree/master/mcgeachy-2018/figure-3.

### Mutational analysis of *RPL28*

#### Genomic RPL28 deletion.

BY4741 yeast were transformed with plasmid pNTI592 (pFA6a ARS/CEN *RPL28(KpnI) CaURA3*) by standard lithium acetate transformation and transformants were selected on SCD ‐Ura plates. An *rpl28*Δ*::KlLEU2* deletion cassette was amplified from a plasmid bearing the *K. lactis LEU2* marker using primers NI-NI-914 and NI-NI-915 and purified by a DNA Clean & Concentrator-5 (Zymo D4003). Roughly1.7 μg of purified PCR product was transformed into yeast carrying pNTI592 by standard lithium acetate transformation, and transformants were selected on SCD ‐Leu plates (without selection for uracil prototrophs). Candidate transformants were tested for stable uracil prototrophy on SCD ‐Ura and SCD ‐Ura supplemented with Ura (50 mg / liter) 5-FOA (100 μg / ml; Zymo Research F9001-1) plates.

DNA was extracted from phenotypically positive clones along with a parental control (36) and verified by PCR using 1.0 μl extracted DNA as a template. A 2.0-kb fragment across the wild-type *RPL28* locus was amplified using primers NI-NI-905 and NI-NI-908 using Q5 polymerase according to the manufacturer’s protocol, 64 °C annealing and 80 second extension time for 35 cycles. PCR products were analyzed by gel electrophoresis and the loss of the 2.0 kb *RPL28* band was seen in phenotypically positive clones, though the 2.6 kb *rpl28*Δ*::KlLEU2* band was never observed. Junctions were then verified by PCR using GoTaq Flexi Green (Promega M7660) with 2.0 mM Mg^+2^ for 30 cycles with 50 second extension times. Primer NI-NI-905 was used with primer NI-NI-901 or NI-NI-894 at a 50°C annealing temperature, and primer NI-NI-908 was used with primer NI-NI-912 or NI-NI-899 at a 52 °C annealing temperature. PCR products were analyzed by gel electrophoresis, and deletion candidates showed a 351 bp product from NI-NI-905 and NI-NI-901 whereas wild-type controls showed a 260 bp product from NI-NI-905 and NI-NI-894. Likewise, deletion candidates showed a 744 bp product from NI-NI-908 and NI-NI-912 whereas wild-type controls showed a 391 bp product from NI-NI-908 and NI-NI-899. One positive isolate was selected as NIY414.

#### Mutagenesis and barcoding.

Exon 2 of *RPL28* was amplified by mutagenic PCR from pNTI592 using primers NI-NI-886 and NI-NI-913 in two 16. μl reactions using the GeneMorph II Random Mutagenesis Kit (Agilent 200550) according to the manufacturer’s protocol, with either 6.4 ng or 0.4 ng DNA template. Amplification was carried out for 30 cycles with 54 °C annealing and 1 minute extension. Products were purified using a DNA Clean and Concentrator-5 with elution into 18.0 μl and yielded 1.2 ‐ 1.5 μg DNA total. Residual plasmid was eliminated by adding 2.0 μl 10x CutSmart Buffer and 0.5 μl DpnI (NEB R0176S) and digesting 90 minutes at 37 °C followed by heat inactivation for 20 minutes at 80 °C.

Mutagenized inserts were amplified by PCR using Q5 according to the manufacturer’s instructions, with 25. μl reactions using 0.5 μl GeneMorph product as a template, primers NI-NI-952 and NI-NI-953, and 25 cycles with annealing at 62 °C and 20 second extension time. PCR products were purified using a DNA Clean and Concentrate-5 and ∼1.0 μg product was digested in a 50 μl reaction using 0.5 μl NheI-HF (NEB R3131S) and 0.5 μl MfeI-HF (NEB R3589S) for 1 hour at 37 °C and purified by DNA Clean and Concentrator-5 with elution into 6.0 μl.

Plasmid pNTI619 (pFA6a ARS/CEN *RPL28exon1-(MfeI / NheI) SpHIS5*) was used as a vector, and 2.2 μg of DNA was digested with NheI-HF and MfeI-HF as described above, with the addition of 0.5 μl rSAP (NEB M0371S).

A library was constructed by combining 430 ng of digested, purified vector with 90 ng each of the two mutagenic PCR inserts in a 60 μl ligation using the Quick Ligation Kit (NEB M2200S) according to the manufacturer’s protocol. Water was added to 150 μl, and ligation products were purified using a DNA Clean and Concentrator-5 with elution into 6.0 μl. The ligation was transformed into 3 x 100 μl aliquots of XL1-Blue chemically competent cells, with 2.0 μl purified ligation product per transformation, according to the manufacturer’s protocol. The transformations were pooled and a small aliquot was diluted and plated on LB Carb plates while the rest of the transformation was used to inoculate a turbidostat containing LB Carb and grown for 20 hours at 37 °C at an OD of ∼1.4, and a density of ∼7.5e8 colony forming units / ml as assessed by plating dilutions on LB Carb plates. Bacteria from 135 ml of culture were used to harvest plasmid DNA using a Qiagen HiSpeed Midiprep (Qiagen 12643) eluted into 1.0 ml according to the manufacturer’s instructions, yielding 88 μg of library DNA.

#### Barcode-to-variant assignment.

A “tagmentation” reaction was carried out using the Nextera XT Library Preparation Kit (Illumina FC-131-1024) according to the manufacturer’s instructions, except that in the library amplification, primers NI-NI-957 and NI-NI-958 were used in place of the Index adapters. The reactions were purified using 0.6 volumes of AMpure XP beads to select for larger inserts. A second round of amplification was performed in two 50 μl PCRs using Q5 according to the manufacturer’s instructions, using primers NI-NI-956 and NI-NI-957 with 1.0 μl of tagmentation library as an input to carry out 10 cycles of amplification at 67 °C annealing and 30 second extension. PCR products were purified using AMpure XP beads, one with 30 μl (0.6 volumes) and one with 50 μl (1.0 volumes) of beads. Libraries were analyzed by TapeStation and combined with the intention of producing equimolar coverage of 300 bp to 850 bp fragments. Libraries were then subjected to asymmetric, paired-end sequencing on the Illumina MiSeq to read 25 bp from read 1 and 125 bp from read 2.

Sequencing data were analyzed by trimming the constant sequence from the end of the read 1 barcode and then renaming read 2 sequences based on the read 1 barcode to facilitate grouping together all reads from the same barcode. Renamed read 2 sequences were aligned to a plasmid reference sequence using Bowtie2 (33) and then grouped according to read name / barcode using SAMtools (34). Read 2 sequences were then collected to determine a consensus sequence from the intron branch point through the end of the coding sequence (bases 1097 through 1546 inclusive). Barcodes missing more than 2 nucleotides or showing more than 1 heterogeneous position (i.e., a position with multiple reads supporting each of two or more variant calls) were excluded from further analysis; a substantial number of barcodes lacked complete sequencing coverage of the mutagenized region. Custom scripts and Rust programs implementing this workflow are available at https://github.com/ingolia-lab/turbidostat/tree/master/mcgeachy-2018 in the “figure-56” and “cyh2” subdirectories.

#### Yeast transformation and selective growth.

The mutagenized *RPL28* library was transformed into NIY414 in eight standard lithium acetate transformations with 3 μg library DNA per transformation. Transformants were inoculated into a turbidostat with 200 ml SCD ‐His at a density of roughly 4e6 cells / ml and grown at a target density of ∼6.5e6 cells / ml. Cells were switched to SCD ‐His supplemented with Ura (25 mg / l added) and 5-FOA (100 μg / ml) and then to SCD ‐His with 200 nM cycloheximide.

Samples were collected from the turbidostat outflow for 15 ‐ 30 minutes. Genotype distributions were assessed by performing two 100-fold serial dilutions and plating 100 μl of this 1:10,000 dilution on YEPD plates. After two days of growth at 30 °C, individual colonies were patched onto SCD ‐His and SCD ‐Ura plates, and auxotrophic markers were scored after two additional days of growth at 30 °C. For all samples, cells were pelleted 5 minutes at 3,000 x g, room temperature, washed in 2.0 ml sterile deionized water, split into two x 1.0 ml samples, pelleted 30 second at 13,000 x g, and the supernatant was removed. Cell pellets were stored at −80 °C.

#### Plasmid recovery and library construction.

Plasmid DNA was extracted from frozen cell pellets using a Zymoprep Yeast Plasmid Miniprep II kit (Zymo Research) according to the manufacturer’s protocol and eluted in 10.0 μl. First round amplification was carried out using 2x Q5 Master Mix with 2.5 μl of mini-prep (or input library at 88 pg / μl) as a template and primers NI-NI-945 and NI-NI-956 with 20 cycles of amplification, 65 °C annealing, and 5 second extension. PCR products were purified from 20 μl PCR by adding 30 μl water followed by 100 μl (2 volumes) AMpure XP beads and purifying DNA according to the manufacturer’s instructions, with elution into 20.0 μl. Second round amplification was carried out using 2x Q5 Master Mix with 1.0 μl purified first-round product as template, along with primer NINI-798 and an index primer (NI-NI-951 or NI-NI-999 through NI-NI-1004) with 12 cycles of amplification, 68 °C annealing, and 5 second extension. PCR products were purified from 20 μl PCR by adding 30 μl water followed by 100 μl (2 volumes) AMpure XP beads and purifying DNA according to the manufacturer’s instructions, with elution into 20.0 μl. Libraries were sequenced on an Illumina HiSeq 4000.

#### Barcode analysis.

Constant sequences were removed from the ends of sequencing data using Cutadapt and distinct barcodes were counted in each sample and tabulated. Barcodes without at least 3 reads counted in the input library sample were discarded. Raw barcode counts were used to compute Spearman rank correlations between samples, which were then used to produce a distance matrix for hierarchical clustering in R. Custom scripts and Rust programs implementing this workflow are available at https://github.com/ingolia-lab/turbidostat/tree/master/mcgeachy-2018 in the “figure-4def” and “cyh2” subdirectories.

#### Variant phenotype analysis.

Barcodes with complete variant sequence information were selected and abundance changes were analyzed using DESeq2. Barcode fitness in normal growth was assessed by treating the post-transformation samples “His æ1 (A)” and “His æ2 (B)” as replicates, the counter-selected samples “5FOA æ1 (C)” and “5FOA æ2 (D)” as replicates, and assessing the fold-change post-5FOA relative to pre-5FOA. Barcode fitness in cycloheximide was assessed by comparing the two counter-selected samples treated as replicates against the final cycloheximide-treated sample “Chx æ2 (F)” with no replication.

The fitness of single-nucleotide variants was assessed by selecting all single-nucleotide variants represented by at least three distinct barcodes and taking the median fitness across all barcodes linked to the variant. The minimum fitness observed in variants expected to be silent (synonymous substitutions or intronic mutations not affecting key splicing signals) were used to define the lowest “healthy” fitness. The maximum fitness observed in variants expected to be non-functional (nonsense mutations and frameshift mutations) were used to define the highest “dead” fitness. Fitnesses between these two levels were defined as “sick”.

Single residue missense variants were assessed by selecting all substitutions represented by at least three distinct barcodes and taking the median fitness across all barcodes linked to the substitution. Position-specific scoring matrix effects were determined from the PTZ00160 PSSM. Cycloheximide resistance was analyzed only for amino acid substitutions classified as “sick” or “healthy” after 5-FOA counterselection; those classified as “dead” were excluded from further analysis. The maximum cycloheximide fitness observed in presumptively silent nucleotide variants as described above was used to define the threshold for cycloheximide sensitivity. Custom R scripts and Pymol commands implementing this workflow are available at at https://github.com/ingolia-lab/turbidostat/tree/master/mcgeachy-2018 in the “figure-56” subdirectory.

#### Cycloheximide sensitivity.

Individual cycloheximide resistant mutations, along with wild-type *RPL28*, were reconstructed in pNTI619 using standard molecular cloning techniques and verified by Sanger sequencing. Plasmids were transformed into NIY414 by lithium acetate transformation and clonal transformants were isolated on SCD ‐His. Transformants were re-streaked onto SCD ‐His +5-FOA plates in order to isolate clones lacking Ura-marked pNTI592.

Overnight cultures were grown in YEPD at 30 °C with shaking and then diluted to an A_600_ of 0.1. YEPD media containing cycloheximide at various concentrations (40, 80, 160, 320, and 640 nM) was used to prepare a 96-well clear round-bottom plate (Corning 353077) and an equal volume of diluted cells were added to achieve a final A_600_ of 0.05 and a final cycloheximide concentration of 0, 20, 40, 80, 160, or 320 nM. The plate was sealed with a gas-permeable membrane (Diversified Biotech, BERM-2000) and cells were grown in a Tecan Infinite M1000 plate reader at 30 °C with shaking. Absorbance was measured every 10 minutes for 16 hours. Growth rates were determined by fitting absorbance data using the grofit package (37).

## ACKNOWLEDGEMENTS

We thank members of the Ingolia lab and L. Lareau for discussion. We also thank H. T. Wong, D. Tullman-Ercek, A. Ollodart, M. Dunham, E. Tung, and W. Luddington for discussions and use of the turbidostats. This work was supported by the Searle Scholars Program 11-SSP-229 (N. T. I.) and an NIH New Innovator’s Award DP2 CA195768-01 (N. T. I.). This work used the Vincent J. Coates Genomics Sequencing Laboratory at UC Berkeley, supported by NIH Instrumentation Grants S10 RR029668, S10 RR027303, and S10 OD018174.

## AUTHOR CONTRIBUTIONS

A.M. and N.I. constructed turbidostats and carried out growth experiments based on a design by N.I. A.M. constructed, sequenced, and analyzed expression libraries. N.I. designed and analyzed *RPL28* variant libraries. Z.M. constructed barcode sequencing libraries and tested cycloheximide resistance of individual variants.

## COMPETING INTERESTS

The authors declare no competing financial interests.

